# Targeted gene correction and functional recovery in achondroplasia patient-derived iPSCs

**DOI:** 10.1101/801415

**Authors:** Huan Zou, Mingfeng Guan, Yundong Li, Fang Luo, Wenyuan Wang, Yiren Qin

## Abstract

**Background:** Achondroplasia (ACH) is the most common genetic form of dwarfism and belongs to dominant monogenic disorder caused by a gain-of-function point mutation in the transmembrane region of *FGFR3*. There are no effective treatments for ACH. Stem cells and gene-editing technology provide us with effective methods and ideas for ACH research and treatment.

**Methods:** We generated non-integrated iPSCs from an ACH girl’s skin and an ACH boy’s urine by Sendai virus. The mutation of ACH iPSCs was precisely corrected by CRISPR-Cas9.

**Results:** Chondrogenic differentiation ability of ACH iPSCs was confined compared with that of healthy iPSCs. Chondrogenic differentiation ability of corrected ACH iPSCs could be restored. These corrected iPSCs displayed pluripotency, maintained normal karyotype, and demonstrated none of off-target indels.

**Conclusions:** This study may provide an important theoretical and experimental basis for the ACH research and treatment.

## Background

Achondroplasia (ACH), the most common genetic form of short-limb dwarfism, is an autosomal dominant monogenic disorder (MGD) caused by a gain-of-function point mutation in the transmembrane region of fibroblast growth factor receptor 3 (*FGFR3*). Currently there are two mutation sites reported - Gly380Arg and Gly375Cys, and the former occupies a vast majority of ACH patients [1, 2]. Because homozygous ACH patients show much more severe symptoms and rarely survive [3, 4], almost all survivors are heterozygous patients. The estimated frequency of ACH is 1 in 25,000 (Male 1/15000), with at least 80% of the cases being sporadic [5]. The clinical symptoms of achondroplasia are evident at birth. The typical physical traits are mainly proximal shortening of the extremities, genu varum, trident hand, limitation of elbow extension, exaggerated lumbar lordosis, megalencephaly, and characteristic facies with frontal bossing. Symptoms of radiological diagnosis include small cuboid vertebral bodies with progressive narrowing of the caudal interpedicular distance, lumbar lordosis, thoracolumbar kyphosis with occasional anterior beaking of the first and second lumbar vertebrae, small iliac wings with a narrow greater sciatic notch, and short tubular bones with metaphyseal flare and cupping [6]. Complications include sleep apnea and recurrent ear infections [1]. Like many other MGDs, there are no effective treatments for ACH even though the mutant gene has been identified for many years [1]. Current therapeutic methods for ACH mainly include limb lengthening for short stature and the treatment of some clinical complications, such as ventricular shunts for hydrocephalus, and decompression surgery for spinal cord compression [7]. Though there are many study about pharmacotherapy, such as growth hormone [8], C-type natriuretic peptide analog [9] and statin [10], which might possess potential to improve the symptoms of ACH, the safety and efficacy need to be further explored and confirmed. Fortunately, stem cell research provides potential treatments for ACH. Patient-derived stem cells can aid scientists in the investigation of specific molecular mechanisms and in the discovery of new drugs [10, 11]. In addition, via powerful genome editing tools, such as clustered regulatory interspaced short palindromic repeat (CRISPR)-Cas9 system [12], mutation of ACH stem cells can be corrected *in vitro*. Through safety assessment in animal models *in vivo*, stem cell transplantation may provide a novel therapy [11]. For example, recently Perlingeiro and his colleagues used limb girdle muscular dystrophy type 2A patient-derived iPSCs which were caused by mutations in the Calpain 3 (CAPN3) to perform gene correction by CRIPR-Cas9, and transplanted gene-corrected myogenic progenitors into a mouse model that combined immunodeficiency with a lack of CAPN3. They found that the transplantation corroborated the rescue of CAPN3 mRNA [13].

In this study, we isolated and cultured skin, urine, and white adipose-derived somatic cells from three ACH patients with Gly380Arg mutation, including a girl, a boy, and an adult male. Further we generated non-integrated iPSCs from the ACH girl’s skin and the ACH boy’s urine. However adipose-derived mesenchymal stem cells (AD-MSCs) from the adult male could not be reprogrammed to iPSCs. We found that the chondrogenic differentiation ability of ACH iPSCs was confined compared with that of healthy iPSCs. When the mutation of ACH iPSCs was precisely corrected by CRISPR-Cas9, their chondrogenic differentiation ability was restored. In addition, these corrected iPSCs kept pluripotency and maintained normal chromosomal number and structure. Our study may provide an important theoretical and experimental basis for the ACH research and treatment.

## Methods

### Research subjects and ethics statement

Research subjects in this study included two children and one adult male. Skin and urine samples were obtained from the children. Liposuction was performed on the adult male. All human subject protocols were reviewed and approved by the Ethical Review Board of the Renji Hospital, Shanghai (License Number 2014102805). All subjects signed the informed consent.

### Isolation and culture of skin fibroblasts (SFs)

Skin biopsies were performed on the locally anesthetized lower legs by a sterile 3 mm skin punch. The skin tissues were cut into smaller pieces and placed in a 6-well plate, and grew for 2 weeks in the fibroblast medium that included DMEM, 10% fetal bovine serum (FBS), penicillin/streptomycin (P/S) and glutamine. Above regents were all from ThermoFisher.

### Culture of urine-derived cells (UCs)

We followed the method established by Pei and his colleagues [14] for the culture of UCs. Briefly, urine samples were collected into 50 ml tubes and centrifuged at 400 g for 10 minutes. The supernatants were sucked out carefully, with only about 1 ml at the bottom kept in the tube. The remaining, after being resuspended by PBS which contained amphotericin B (Selleck) and P/S, were centrifuged again. Thereafter, with the supernatants discarded, the pellets were suspended and cultured into 12-well plates by the primary medium that contained DMEM, Ham’s F-12 (ThermoFisher), 10% FBS, REGM Renal Epithelial Cell Growth Medium SingleQuots Kit (Lonza), amphotericin B and P/S. After 4 days, the medium was carefully changed to proliferation medium (Lonza REBM BulletKit).

### Isolation and culture of AD-MSCs

Adipose biopsy was carried out by liposuction. Isolation and culture of AD-MSCs were performed according to our previous report [15]. Briefly, adipose tissues were digested by 0.1% type I collagenase (Sigma) in PBS on a shaker at 37°C for 1 hour. Cell suspensions were then centrifuged and the pellet at the bottom was stromal vascular fraction (SVF). Finally, the SVF was suspended and cultured in the fibroblast medium.

### PCR and Sanger Sequencing

Total genomic DNA were extracted from patient-derived peripheral blood, SFs, UCs, AD-MSCs and iPSCs by using the Genomic DNA Extraction Kit (TaKaRa). PCR was performed by using Q5 High-Fidelity DNA Polymerase (New England Biolabs, NEB). *FGFR3* primers were:

Forward: 5’-AGGAGCTGGTGGAGGCTGA-3’,

Reverse: 5’-GGAGATCTTGTGCACGGTGG-3’.

PCR reactions were then purified by using GeneJET Gel Extraction Kit (ThermoFisher) and sequenced by the Genewiz.

### Generation of non-integrated iPSCs

Inductions of iPSCs from SFs, UCs and AD-MSCs were performed by CytoTune iPS 2.0 Sendai Reprogramming Kit (ThermoFisher). Briefly, approximately 200,000 donor cells were cultured in a 6-well plate and transfected with the Sendai virus two days later. Twenty-four hours after transfection, the media for somatic cells were replaced with fresh ones every other day. On day 8 after transfection, the cells were digested by trypsin and 5000 cells were seeded into a 6-well plate to culture. Thereafter, the media were replaced by E8 (StemCell) every day. Three to four weeks after transfection, single cell colonies were picked up for expansion and characterization.

### Chondrogenic differentiation of iPSCs

This method referred to a report of Tsumaki laboratory^7^. When iPSCs formed high-density cell colonies in E8, the media were changed to the initial chondrogenic medium that included DMEM/F12 (ThermoFisher), 10 ng/ml of Wnt3A (R&D), 10 ng/ml of Activin A (Peprotech), 1% ITS (ThermoFisher) and 1% FBS (ThermoFisher). On day 3 after the chondrogenic induction, the medium was changed to the basal medium that contained DMEM (ThermoFisher), 1% ITS, 1% FBS, 2 mM L-glutamine, non-essential amino acids, 1mM Napyruvate (ThermoFisher), 50 μg/ml ascorbic acid (Sigma), 10 ng/ml BMP2 (Peprotech), 10 ng/ml TGFβ1 (Peprotech) and 10 ng/ml GDF5 (Peprotech). From day 3 to day 14, 10 ng/ml bFGF (Peprotech) was added to the chondrogenic medium. After Day 42, the medium was changed to the fibroblast medium.

### Alkaline phosphatase (AP) staining

AP staining was performed by Vector Blue AP Substrate Kit (Vector). The procedure was conducted in accordance with the instruction.

### Gene correction of ACH-iPSCs

sgRNAs were designed by using the Guide Design Resources. Then these sgRNAs were annealed and ligated to the pSpCas9(BB)-2A-RFP plasmid which was digested by Bbs I (NEB) enzyme. One million iPSCs were used for transfection targeting plasmid and ssODNs. The cells were electroporated by using the Human Stem Cell Nucleofector Kit 2 (Lonza) and the Nucleofector 2b Device (Lonza). Twenty-four to forty-eight hours after electroporation, about 5000 RFP positive cells were re-seeded into a 100 mm plate by FACS (BD Aria II). One week later, single cell colonies were picked up and expanded for sequencing analysis. Specific procedure is in Supplementary material and method.

### Off-target effect analysis

Eighteen potential “off-target” sites of the sgRNA used in this research were given by the Guide Design Resources. PCR products of them were sequenced for off-target effect analysis. The sequences of the primers for off-target sites were listed in supplementary Table S1.

### Cell immunofluorescence

iPSCs were fixed in 4% paraformaldehyde solution for 15 minutes at room temperature (RT). Then they were permeabilized by using 0.1% Triton X-100 in PBS for 20 minutes at RT. After they were blocked in 5% goat serum for 1 hour, the cells were incubated overnight at 4°C by primary antibodies, including Nanog (Abcam), SSEA-1 (Invitrogen), Oct4 (Cell signal), SSEA-4 (ThermoFisher), Sox2 (Epitomics) and TRA-1-60 (Abcam). Finally, these cells were incubated by fluorescently coupled secondary antibodies (Invitrogen) and 4’,6-diamidino-2-phenylindole (DAPI; Sigma) for 1 hour at RT. Images were captured on a confocal microscope (Leica).

### Karyotyping

iPSCs were cultured in E8 with 0.1 μg/ml colchicine (ThermoFisher) for 2 hours. After the medium being removed, they were incubated in 0.56% potassium chloride for 40 minutes, and fixed in methanol and acetic acid (3:1 in volume) overnight. The cell suspensions were dropped onto cool slides and stained with Giemsa (ThermoFisher) for 15 minutes. More than 15 metaphase spreads of each sample were analyzed.

### RNA extraction and RT-qPCR

Total RNAs were extracted from cells by using TRIzol RNA Isolation Reagents (Thermofisher). cDNAs were synthesized from total RNA with a Hifair II 1st Strand cDNA Synthesis Kit (Yeasen) for qPCR. The qPCR procedure was carried out according to the kit instructions (TB Green Premix Ex Taq kit, TakaRa). Primers used were as follows:

*SOX9* F: AGACCTTTGGGCTGCCTTAT

R: TAGCCTCCCTCACTCCAAGA

*COL2A1* F: TTTCCCAGGTCAAGATGGTC

R: CTTCAGCACCTGTCTCACCA

*ACAN* F: AGGCAGCGTGATCCTTACC

R: GGCCTCTCCAGTCTCATTCTC

*β-ACTIN* F: TGGCACCACACCTTCTACAATGAGC

R: GCACAGCTTCTCCTTAATGTCACGC

### Safranin O staining

Safranin O staining Kit (ScienCell Research Laboratories, Inc. #8384) was used to perform this experiment. The procedure was in accordance with the instruction.

### Statistical analysis

Data are expressed as the mean ± standard error of the mean (SEM) after comparison by t-test. Significant differences between groups were assessed by Prism software. Differences with P < 0.05 were considered statistically significant.

## Results

### Identification, isolation and culture of somatic cells from ACH patients

We recruited three ACH patients, including an 8-year-old girl, a 7-year-old boy and a 37-year-old adult male. Genetic mutations of the donors were confirmed by Sanger sequencing, and they are all heterozygous mutations of Gly380Arg (Fig. 1a). In addition, their clinical symptoms are consistent with the NIH-defined characteristics, such as short arms and legs, enlarged head, and prominent forehead. We punched skin tissue and collected urine from the children to culture SFs and UCs. Via liposuction, we obtained adipose tissue from the adult male to culture AD-MSCs. After the small pieces of skins were placed and cultured in the dish for 2 weeks, SFs gradually climbed out to proliferate (Fig. 1b, c). After the urine-derived pellets were cultured for one week, UCs formed colonies and showed epithelial cell morphology (Fig.1d, e). After the SVF from adipose tissue was seeded and cultured for 2-3 days, AD-MSCs grew out and exhibited a typical fibroblast-like morphology (Fig. 1f, g).

**FIG. 1.**
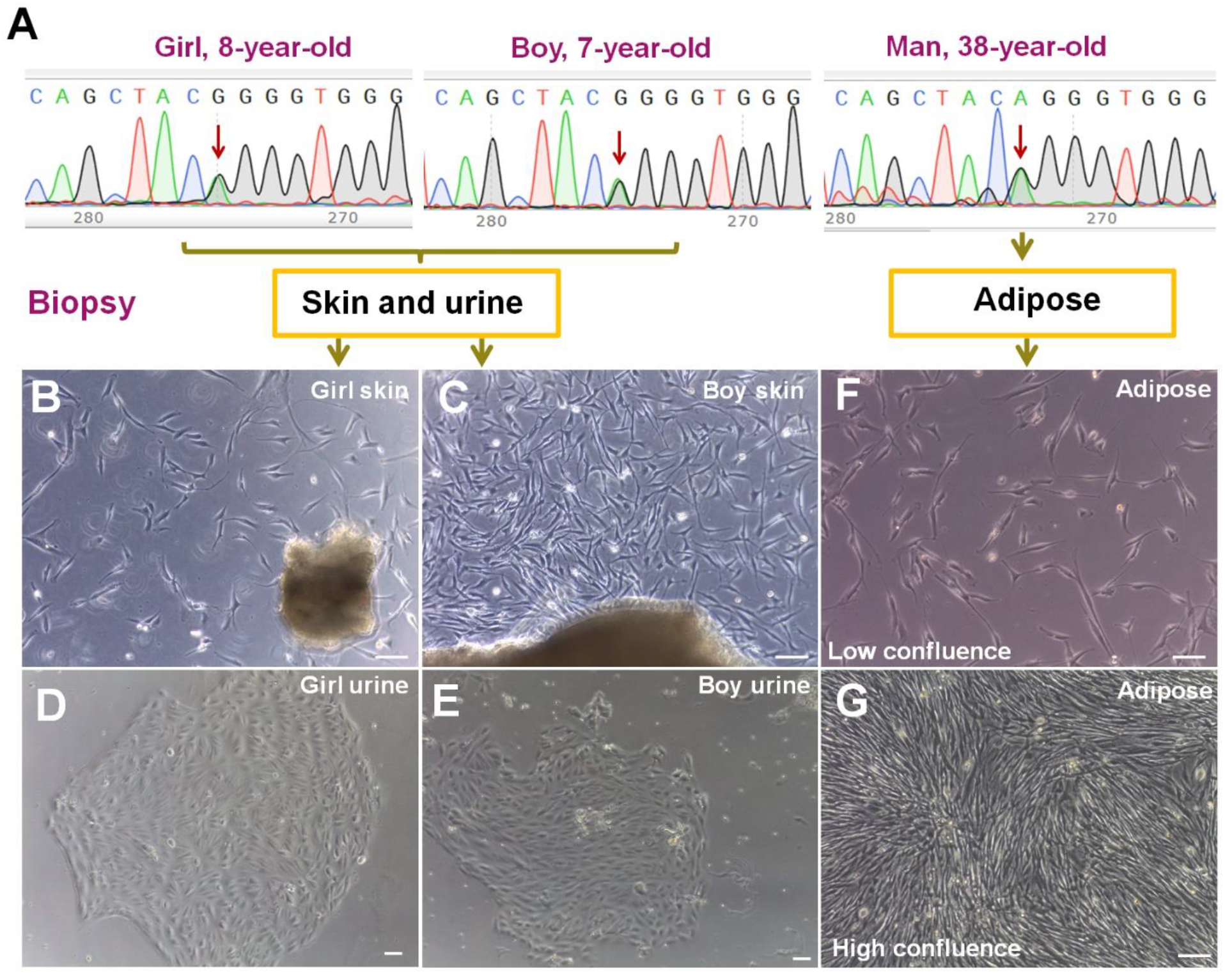
Identification, isolation and culture of somatic cells from ACH patients. **(A)** DNA sequences of the three ACH patients showed that these disorders were all caused by heterozygous mutations of Gly380Arg in the *FGFR3*. **(B, C)** SFs from the girl and the boy. **(D, E)** UC colonies from the girl and the boy showed epithelial cell morphology. **(F, G)** AD-MSCs from the adult male exhibited a typical fibroblast-like morphology. Bar in all panels: 10 μm.

### Generation of non-integrated iPSCs from ACH patients

To generate non-integrated iPSCs from ACH patients, we used Sendai virus to transfect somatic cells. Three weeks after transfection of SFs and UCs, lots of ES cell-like colonies appeared (Fig. 2a, b). After single colonies were picked up and expanded, iPSC lines from the girl’s skin (GF) and the boy’s urine (BU) were established (Fig. 2c, d). These iPSCs expressed pluripotent proteins, including NANOG, OCT4, SOX2, SSEA-4 and TRA1-60, and did not express SSEA-1 (Fig. 2g). They also stained positive for the AP activity (Fig. 2h). Moreover, karyotyping analysis indicated that these iPSCs maintained normal chromosomal number and structure (Fig. 2i). However, it was surprised that iPSCs could not be generated from AD-MSCs. Although small and unhealthy colonies appeared from AD-MSCs (Fig. 2e), they gradually became apoptotic and eventually died (Fig. 2f).

**FIG. 2.**
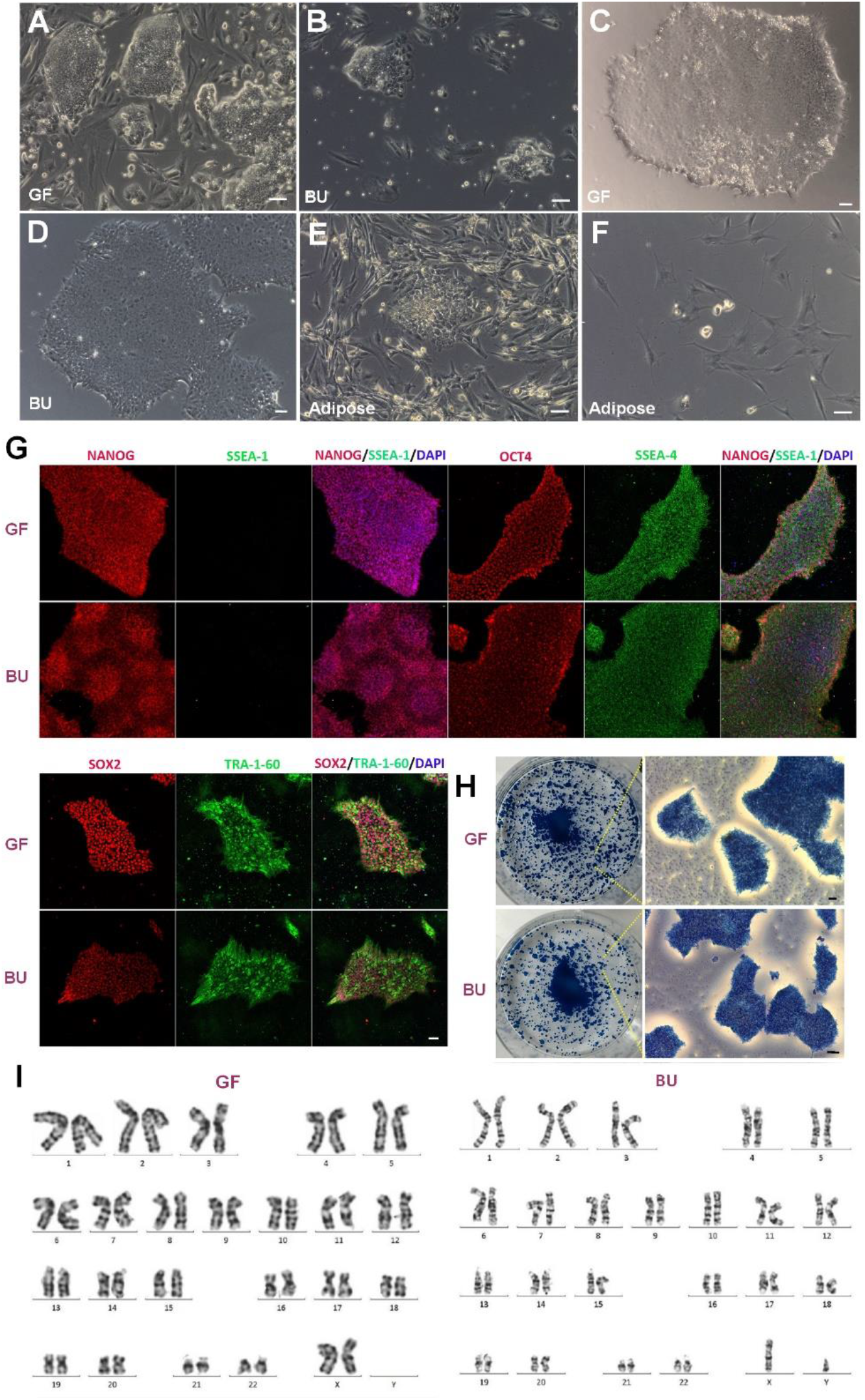
Generation and characterization of non-integrated iPSCs from ACH patients. **(A, B)** 3 weeks ES cell-like colonies from the girl’s SFs (GF) and the boy’s UCs (BU) after transfection. **(C)** Expanded iPSCs from GF. **(D)** Expanded iPSCs from BU. **(E)** 3 weeks ES cell-like colonies from AD-MSCs after transfection were small and unhealthy. **(F)** Colonies from AD-MSCs gradually became apoptotic and eventually died. **(G)** iPSCs from GF and BU expressed pluripotent protein, including NANOG, OCT4, SOX2, SSEA4 and TRA1-60, and did not express SSEA-1. **(H)** iPSCs from GF and BU stained positive for the AP activity. **(I)** iPSCs from GF and BU indicated normal chromosomal number and structure. Bar in all panels: 10 μm.

### Chondrogenic differentiation capability of ACH iPSCs

To confirm whether the mutations would affect the function of ACH iPSCs, we performed the chondrogenic differentiation capability detection. We induced ACH iPSCs and healthy human iPSCs respectively into chondrogenic tissues by a multi-step method (Fig. 3a). We found that the chondrogenic clusters from ACH iPSCs were less and smaller than those from healthy iPSCs (Fig. 3b). By Safranin O staining, we discovered that the cartilage tissues derived from ACH iPSCs showed fewer positive areas and lower cartilage density than those from healthy iPSCs (Fig. 3c). These results suggested that the cartilage tissues derived from ACH iPSCs could produce less cartilaginous extracellular matrix. RT-qPCR results also exhibited that cartilage tissues derived from ACH iPSCs expressed lower chondrocyte-specific genes than those from healthy iPSCs, including *SOX9, COL2A1* and *ACAN* (Fig. 3d). Our results verified that the chondrogenic differentiation capability of ACH iPSCs was confined compared with that of healthy iPSCs.

**FIG. 3.**
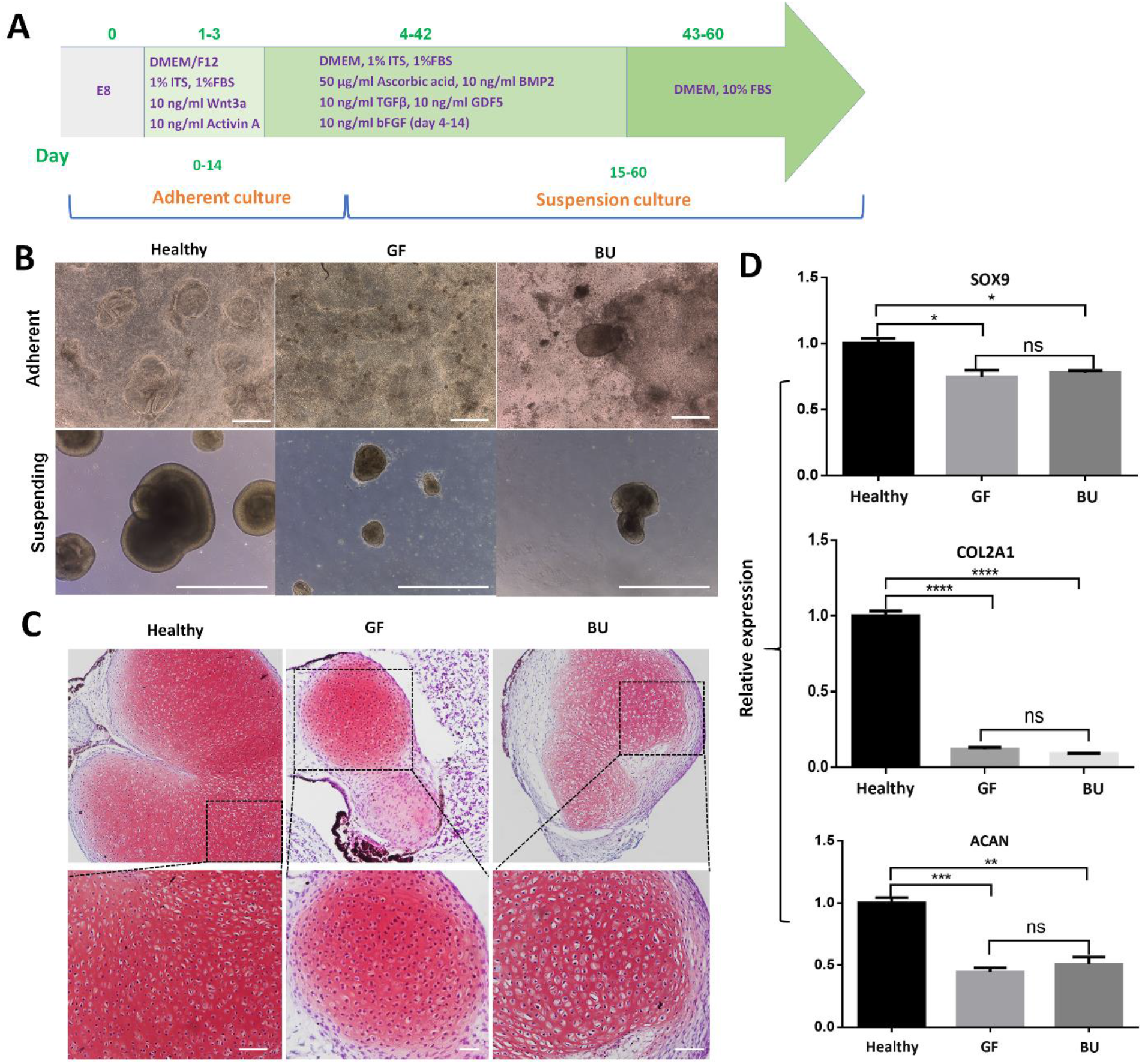
Chondrogenic differentiation of ACH iPSCs. **(A)** A multi-step chondrogenic induction method. **(B)** Chondrogenic clusters from ACH iPSCs were less and smaller than those from healthy iPSCs. Bar: 100 μm. **(C)** Safranin O staining displayed that there were fewer positive areas and lower cartilage density in cartilage tissues from ACH iPSCs than those from healthy iPSCs. Bar: 10 μm. **(D)** RT-qPCR results also exhibited that cartilage tissues derived from ACH iPSCs expressed lower chondrocyte-specific genes, such as *SOX9, COL2A1*, and *ACAN*, than those from healthy iPSCs. Symbol “*” stands for p<0.05. Symbol “**” stands for p<0.01. Symbol “***” stands for p<0.001. Symbol “****” stands for p<0.0001.

### Gene correction of ACH iPSCs by CRISPR-Cas9

To perform the CRISPR-Cas-based gene correction, we designed single guide RNAs (sgRNAs) around the point mutation site of *FGFR3* and a single-stranded oligo DNA nucleotides (ssODNs) homology arm donor which contained 131 nucleotides and the point mutation site (Fig. 4a). We transfected CRISPR-sgRNAs and ssODNs into ACH iPSCs, and 24-48 hours later detected the transfection efficiency by FACS. The ratio of RFP positive cells was 3.6% (Fig. 4b). After analyzing more than 100 RFP positive single cell colonies from skin, we found one completely corrected cell line. Similarly, among more than 100 analyzed RFP positive single cell colonies from urine, we also just found one completely corrected cell line (Fig. 4c). The total efficiency was less than 1%. We found that corrected iPSCs expressed pluripotent proteins, including NANOG, OCT4, SOX2, SSEA-4 and TRA1-60, did not express SSEA-1 (Fig. 4d), and stained positive for the AP activity (Fig. 4e). The karyotyping analysis verified that corrected iPSCs maintained normal chromosomal number and structure (Fig.4f), and the Sanger sequencing showed that there were no off-target indels in them.

**FIG. 4.**
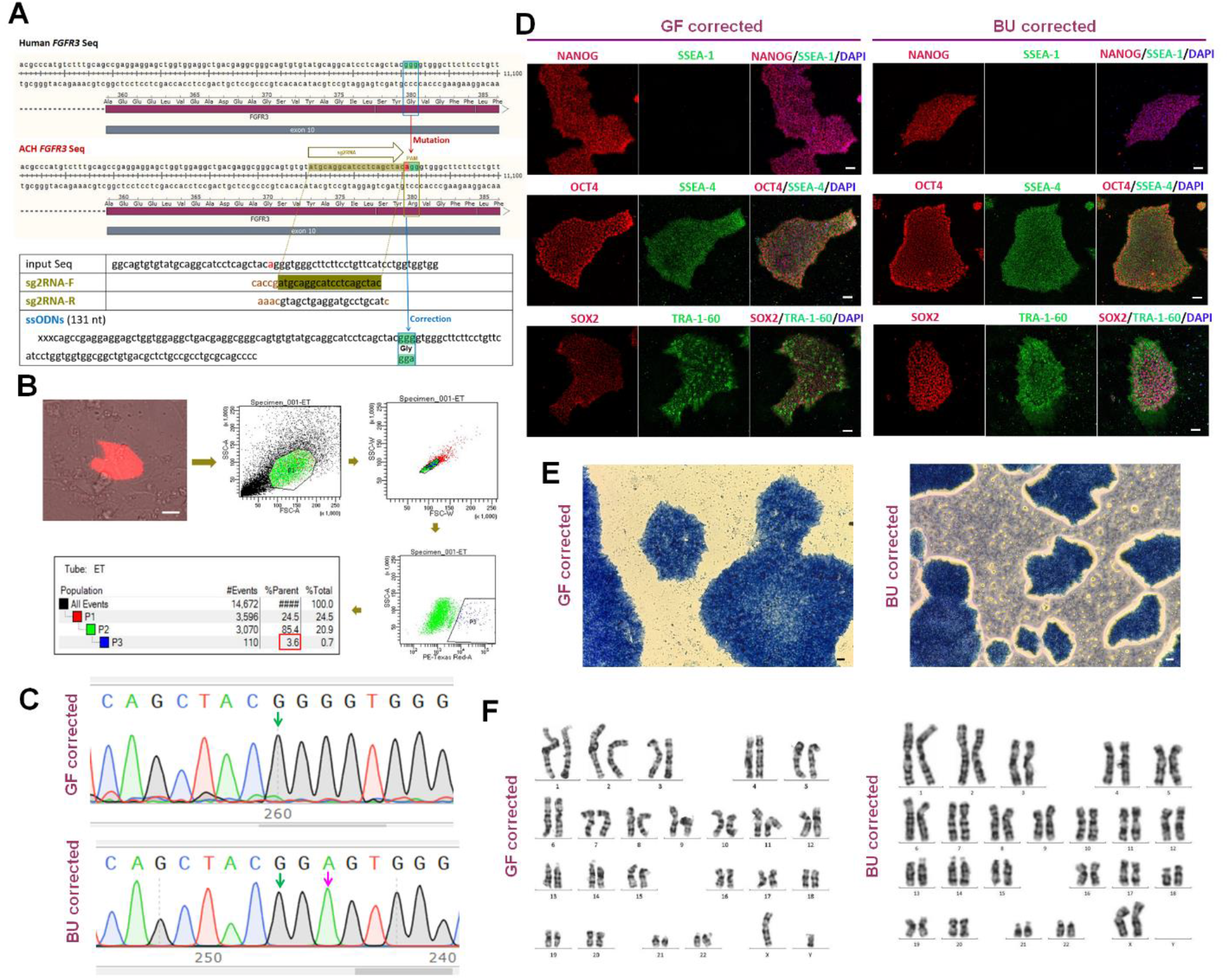
Gene correction of ACH iPSCs by CRISPR-Cas9 and characterization of corrected iPSCs. **(A)** Designed sgRNA and ssODNs around the point mutation site. **(B)** FACS analysis showed that the ratio of RFP positive cells was 3.6%. **(C)** Corrected iPSCs showed normal DNA sequences (Pink arrow indicated a synonymous mutation). **(D)** Corrected iPSCs expressed NANOG, OCT4, SOX2, SSEA4 and TRA1-60, and did not express SSEA-1. **(E)** Corrected iPSCs stained positive for the AP activity. **(F)** Corrected iPSCs showed normal chromosomal number and structure. Bar in all panels: 10 μm.

### Functional recovery of corrected ACH iPSCs

Through morphological observation, we found that the number of chondrogenic clusters from corrected ACH iPSCs obviously increased compared with that from uncorrected cells (Fig. 5a). Via Safranin O staining, we found that there were more positive areas in cartilage tissues from corrected ACH iPSCs than those from uncorrected ones (Fig. 5b). RT-qPCR results also revealed that, compared with cartilage tissues from uncorrected ACH iPSCs, those from corrected ACH iPSCs expressed higher chondrocyte-specific genes - *SOX9, COL2A1*, and *ACAN* (Fig. 5c). These results suggested that the chondrogenic differentiation ability of corrected ACH iPSCs was restored.

**FIG. 5.**
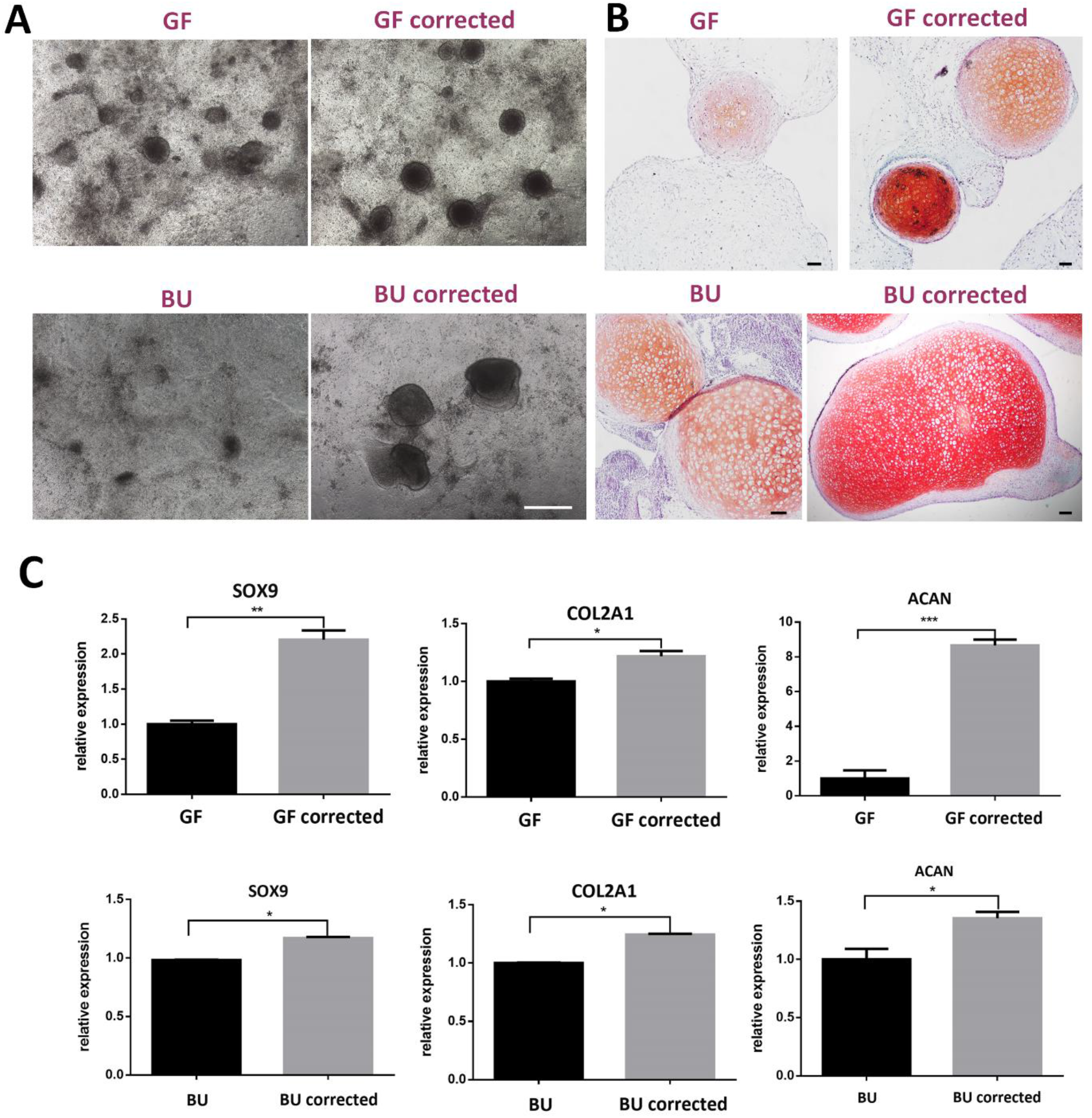
Chondrogenic differentiation of corrected ACH iPSCs. **(A)** The number of chondrogenic clusters from corrected iPSCs obviously increased compared with that from uncorrected cells. Bar: 100 μm. **(B)** Safranin O staining illustrated that cartilage tissues from corrected ACH iPSCs demonstrated more positive areas than those from uncorrected cells. Bar: 10 μm. **(C)** RT-qPCR results also revealed that the expression level of chondrocyte-specific genes - *SOX9, COL2A1*, and *ACAN* from corrected ACH iPSCs were higher than those of ACH iPSCs.

## Discussion

The average life expectancy for an ACH patient was decreased by 15 years compared with that for US population [5]. Moreover, the social concern for this disease is very low. At present there are no effective therapeutic methods for ACH. Fortunately, rapid development of stem cell biology and gene-editing technology provide a promising future for the research and treatment of ACH and other MGDs [11].

In this study we firstly collected three different tissues from ACH patients to culture somatic cells, including the most commonly used skin, the most easily available urine, and a large amount of acquired adipose tissue that can produce multipotent MSCs with chondrogenic differentiation ability. We found that, like skin cells, ACH patient urine-derived cells could be efficiently reprogrammed into iPSCs, which may provide new donor cells for research of MGDs. However, to our surprise, ACH patient-derived AD-MSCs could not be reprogrammed into iPSCs. In fact, previously reported studies [16, 17] and our unpublished results indicated that healthy human AD-MSCs could be reprogrammed to iPSCs more efficiently than skin cells. Given that ACH is a regeneration dysfunction disorder of MSC-derived chondrocyte caused by *FGFR3* mutation, our results suggested that perhaps the point mutation affected the reprogramming ability of AD-MSCs.

Our initial hypothesis was that the point mutation might affect the chondrogenic differentiation ability of ACH iPSCs. Indeed, our experimental results confirmed that the chondrogenic differentiation ability of ACH iPSCs was confined compared with that of healthy iPSCs. When we used the CRISPR-Cas9 technology to correct the mutation of ACH iPSCs, we obtained two completely corrected cell lines, one from skin and the other from urine, among more than two hundred single cell colonies. The efficiency of precise homology directed repair (HDR) was less than 1%. Fortunately, we found that those corrected iPSCs still displayed pluripotency and maintained normal karyotype. The sequencing analysis of potential off-target sites suggested that there were no off-target indels in corrected iPSCs. Finally, we detected whether the function of corrected ACH iPSCs got improved. We found that, via chondrogenic induction and RT-qPCR experiments, the chondrogenic differentiation ability of corrected ACH iPSCs was indeed restored compared with that of uncorrected cells.

## Conclusion

In summary, our study provided an important foundation for the further exploration of ACH research and treatment. At present, we are constructing point mutation mouse models of ACH. In the future, we will transplant corrected ACH patient-derived MSCs or chondrocyte precursor cells into mouse models to verify their function *in vivo* and explore the effect of cell therapy to ACH. Although challenges remain, the clinical application of patient-specific stem cells will be pursued through further advances in basic research (Fig. 6).

**FIG. 6.**
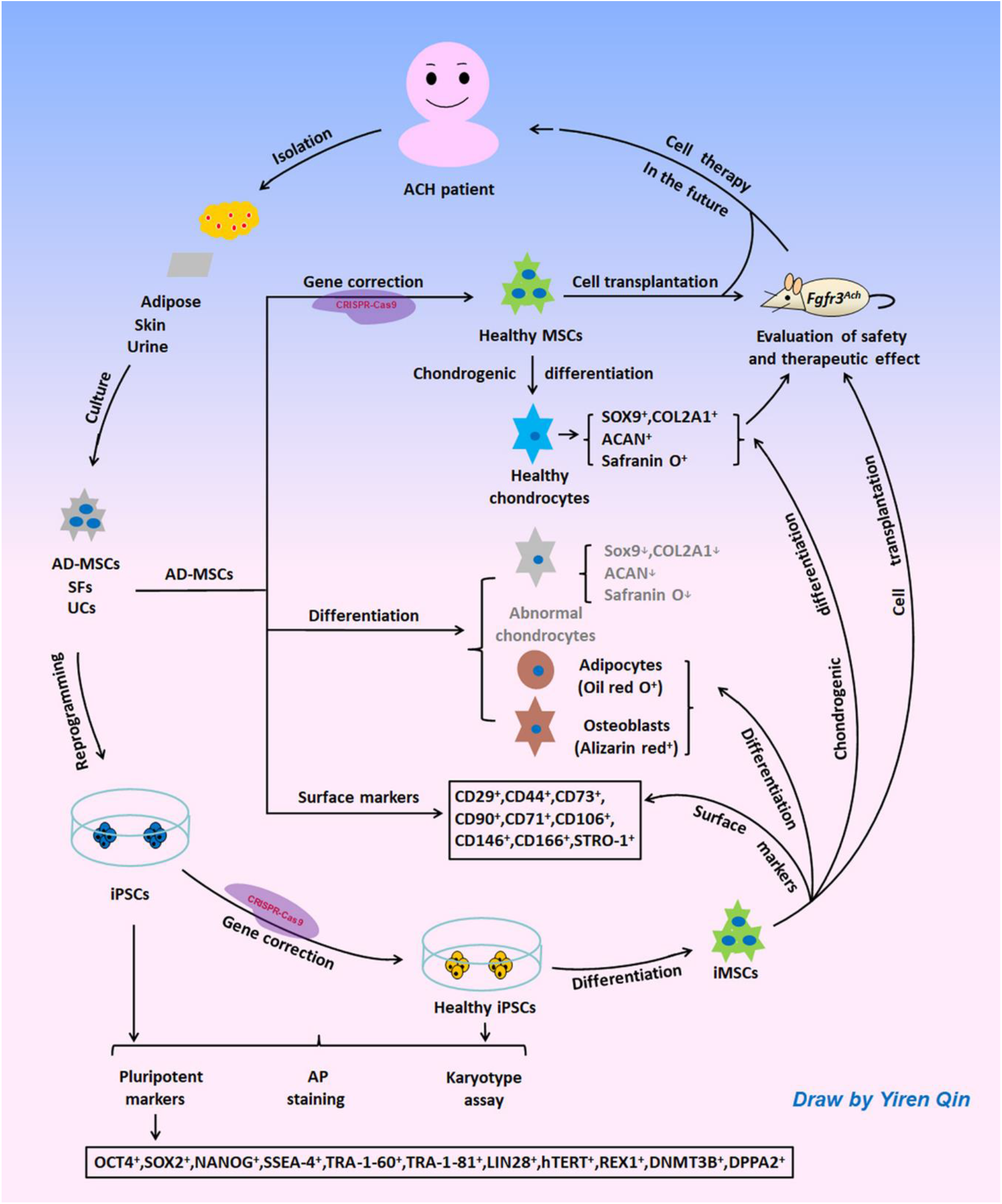
Diagrammatic strategy of ACH stem cell research. Somatic cells can be isolated and cultured from ACH patient-derived adipose, skin, and urine, and can further be reprogrammed into iPSCs. After gene correction of iPSCs or AD-MSCs via CRISPR-Cas9, they can be differentiated into healthy cells, such as MSCs or chondrocyte precursor cells. Then these healthy cells can be transplanted into ACH mouse models to assess their relative safety and therapeutic effects. The clinical application of ACH patient-derived stem cells will be pursued in the future.

## Acknowledgements

We would like to thank Dr. Wei-Qiang Gao for his helpful discussion in conceiving the project, Dr. Hui Yang for providing pSpCas9(BB)-2A-RFP plasmid, Chikai Zhou and Dr. Erwei Zuo for their helpful discussion in designing sgRNA, Dr. Sushan Luo for skin biopsies, Guangrui Cao for her helpful language modification.

## Author contributions

H.Z. and M.G. performed experiments and analyzed data, Y.L., and F.L. performed experiments; W-Y.W. provided experimental platform and financial support; Y.Q conceived and designed the study, performed experiments, analyzed data, provided financial support and wrote the manuscript.

## Funding statement

This work was funded by grants from National Natural Science Foundation of China (General Program 31570992) to Y.Q., National Key R&D Program of China (Grant No. 2018YFA0107903, 2016YFA0501902) and Shanghai Municipal Science and Technology Major Project (Grant No. 2019SHZDZX02) to W-Y.W.

## Availability of data and materials

All data generated or analyzed during this study are included in this published article and its supplementary information file.

## Ethics approval and consent to participate

All human subject protocols were reviewed and approved by the Ethical Review Board of the Renji Hospital, Shanghai. All subjects signed the informed consent.

## Conflicts of Interest

There is no potential conflict of interest.

## Consent for publication

Consent for publication was given by the adult patient and parents of the child patients.

## Supplementary Material

Supplementary data - Ethics statement

Supplemental Table S1

Supplemental material and method

## List of Abbreviations

ACH: Achondroplasia
*FGFR3*: fibroblast growth factor receptor 3
CRISPR: clustered regulatory interspaced short palindromic repeat
MGD: monogenic disorder
CAPN3: Calpain 3
AD-MSCs: adipose-derived mesenchymal stem cells
SFs: skin fibroblasts
UCs: urine-derived cells
FBS: fetal bovine serum
P/S: penicillin/streptomycin
SVF: stromal vascular fraction
AP: alkaline phosphatase
RT: room temperature
DAPI: 6-diamidino-2-phenylindole
GF: girl’s skin
BU: boy’s urine

## Notes

### Competing Interest Statement

The authors have declared no competing interest.

### Summary of Updates

Figure 5, manuscript text

